# Oxidative Stress, Hypoxia and Cellular Metabolism: Unraveling the Effects of Fentanyl on Lung Cancer Cells

**DOI:** 10.1101/2025.07.17.665412

**Authors:** Akanksha Sharma, Rahul K. Das, Andrey Kuzmin, Shobha Shukla, Paras N. Prasad, Supriya D. Mahajan

## Abstract

Fentanyl, a widely used opioid analgesic for cancer pain management, is effective but requires cautious administration due to its potential for respiratory depression. Beyond its analgesic properties, fentanyl’s broader impact on cancer biology and biochemical alterations in lung carcinoma cells remains underexplored. This study investigates fentanyl’s influence on oxidative stress, mitochondrial function, hypoxia-inducing factors, apoptosis, and cytokine production in A549 lung cancer cells. Our findings reveal that fentanyl increases reactive oxygen species (ROS) generation, disrupts cellular homeostasis, induces DNA damage, and alters key signaling pathways, potentially affecting tumor metabolism and progression. Our “Ramanomics” data further highlight fentanyl-driven changes in key ions (inorganic phosphate, calcium) and biomolecules (glycogen, phospholipids, proteins, nucleic acids, and enzymes) at subcellular mitochondrial levels. These insights contribute to understanding fentanyl’s mechanistic impact on lung cancer progression and may inform optimized therapeutic strategies.

**GRAPHICAL ABSTRACT:** 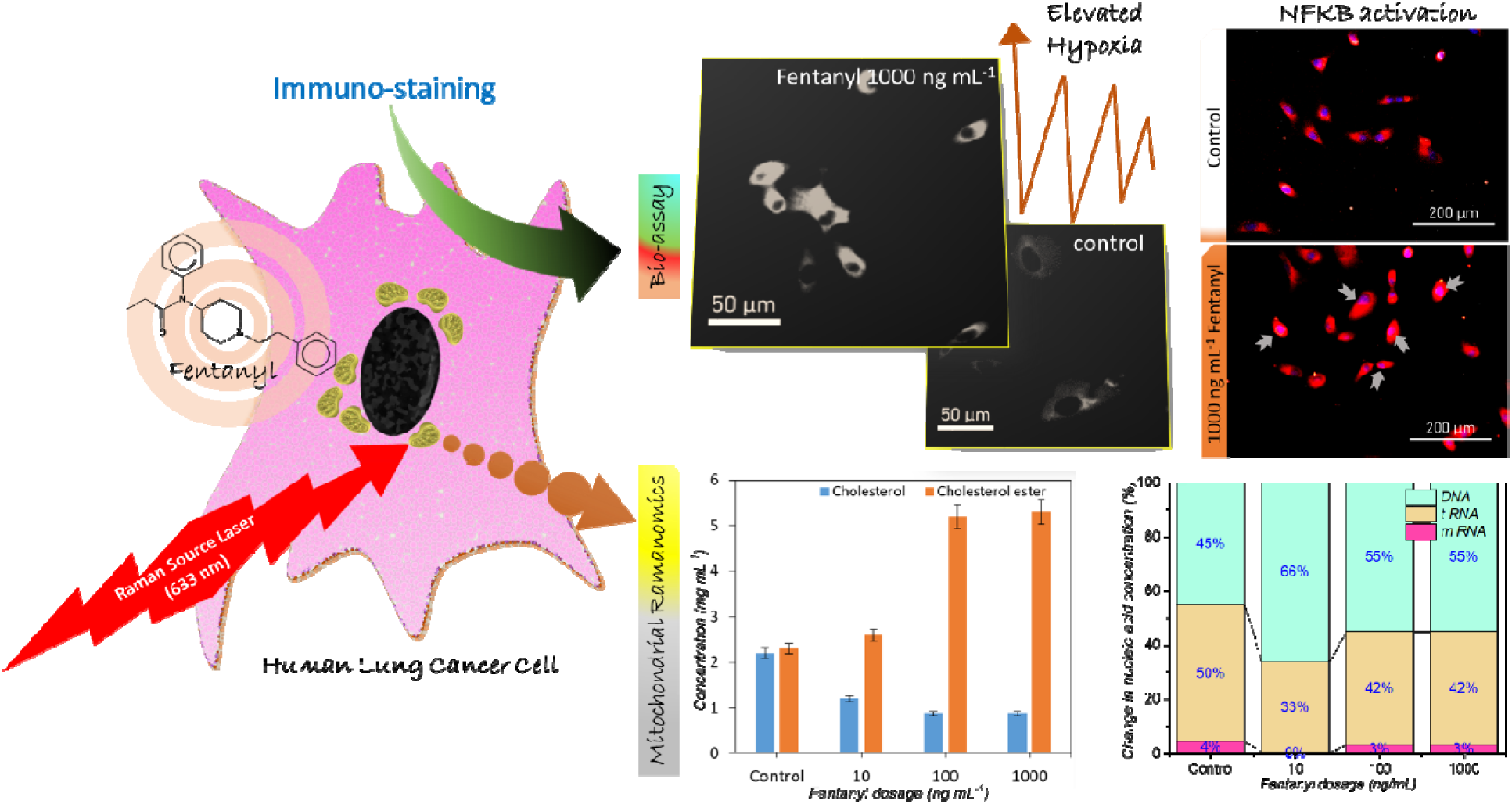

## INTRODUCTION

Lung cancer is a highly malignant carcinoma with an estimated annual mortality of 1.1 million worldwide^1,2^. Fentanyl is an opioid agonist for μ-opioid receptor (MOR), which has been commonly used in clinical anesthesia, and to treat mild to moderate cancer pain^3^. Transdermal fentanyl has FDA approval for patients with moderate to severe chronic non-cancer and cancer-associated pain. Fentanyl transdermal skin patches are used in lung cancer patients and are associated with decreased pain in 60 % of patients upon switched over from oral analgesics. Additionally, adverse effects associated with oral/ injectable opioids such as nausea and constipation, and other issues such as cachexia and mechanical obstruction were significantly reduced^4^. Randomized, controlled, open-label trials suggest that transdermal fentanyl is safe and as effective as sustained-release oral morphine (SROM) in treating chronic cancer pain^5^. Non-small cell lung cancer (NSCLC) accounts for approximately 80% of all patients with lung cancer and therapeutic advances that include radiation, anti-angiogenesis-, and immune therapy, have resulted in improved prognosis of patients with NSCLC, however their five-year survival rate remains below < 20%. In terminally ill lung cancer patients, alleviation of pain is the primary objective and a once-a-day fentanyl transdermal patch is an acceptable long-term strategy for the treatment of cancer-associated pain control in lung cancer patients^6,7^. Fentanyl causes respiratory depression, a potentially lethal side effect with overdose, by acting on μ-opioid receptors in brainstem regions regulating breathing^8^. μ-opioid receptors are present on the NSCLC cell line A549 and MOR is upregulated in lung tissue from patients with NSCLC. Overexpression of MOR is believed to promote tumor growth and metastasis in human NSCLC^8^.

Limited information is available about the characteristics and mechanisms of action of Fentanyl in lung cancer cells. Fentanyl induces respiratory depression by blunting peripheral chemoreceptor sensitivity to CO□ and suppressing central respiratory drive, which may produce systemic hypoxia and thereby alter tumor cell energy metabolism and proliferation. We hypothesize that fentanyl also elevates oxidative stress in lung cancer cells by increasing reactive oxygen species (ROS) production, disrupting redox homeostasis, causing DNA damage, and modulating signaling pathways that regulate growth and survival.

In our study, we investigated the impact of fentanyl on reactive oxygen species (ROS) generation, singlet oxygen production, mitochondrial function, hypoxia-inducible factor (HIF) expression, apoptotic pathway activation and cytokine secretion in A549 lung adenocarcinoma cells, and elucidated the underlying molecular mechanisms. Our results demonstrate that fentanyl modulates redox homeostasis, mitochondrial activity, HIF signaling, apoptotic machinery and inflammatory mediator release, revealing its multifaceted effects on lung cancer cell biology. Given these findings and despite its clinical utility for cancer pain relief, caution is warranted when prescribing transdermal fentanyl patches to patients with lung cancer, as fentanyl may exert deleterious effects on tumor progression.

## MATERIALS AND METHODS

### Cell Culture

The human NSCLC cell line (A549) was purchased from the American Type Culture Collection (ATCC, Manassas, VA, USA). Cells were cultured in RPMI-1640 medium (Gibco, Carlsbad, CA, USA), which were supplemented with 1% antibiotic-antimycotic mixture (Gibco) and 10% fetal bovine serum (FBS; Hyclone, Logan, UT, USA). The cultures were maintained in the atmosphere with humidified 5% CO_2_ and 95% air at 37°C.

### Fentanyl

The fentanyl hydrochloride (CAS 1443-54-5, 1 mg) was procured from Cayman Chemical and dissolved in culture media to prepare 1.0 mg/mL drug concentration. Fentanyl concentrations used in experimental paradigms were 10 – 100ng/mL based on previous reports (Kong L, et al 2021).

### Cell viability assay

Cell viability was analyzed using a Cell Counting Kit-8 according to the manufacturer’s instructions (CK04, Dojindo Molecular Technologies). Briefly, 10,000 A549 cells in 100 μl media/well were added to a 96-well plate. After the cells were attached, Fentanyl at a concentration of (10-1000 ng/ml) in 10 μl volume was added in triplicate, and incubated for 24 hr. Post incubation, 10 μl of CCK-8 solution was added to each well. The cells were then incubated at 37°C for 2 hr. After incubation, the absorption at 450 nm was measured using a Spectrophotometer Microplate Reader. The results were expressed as a percentage of the values of the untreated control set at 100% after subtraction of blank (no cell wells). The cell viability was normalized against untreated control cells and percentage cell viability was calculated using Equation 1:

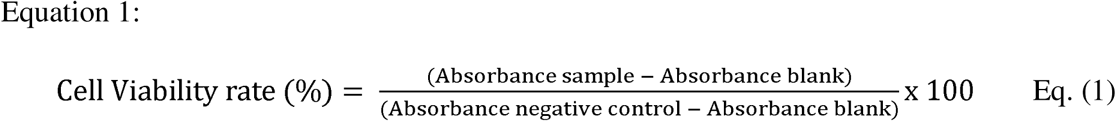

Where, Absorbance negative control= untreated control (Cells + media + CCK-8)

Absorbance sample = Cells + Test drug + Media + CCK-8

Absorbance blank = Media (no cells) + CCK-8

### Caspase 3/7 Detection

For specific staining of dying cells, activated caspases associated with apoptosis, Fentanyl treated A549 cells were visualized with CellEvent(R) Caspase 3/7 Detection Reagent (ThermoFisher Scientific C10423). A549 cells were grown to ∼80% confluence in a 20mm petri dish with a glass bottom (Cat # 801001 NEST), then treated for 24 hr with Fentanyl (10-1000 ng/ml). After incubation, cells were washed with 1X PBS and treated with freshly prepared Caspase 3/7 Detection. (1:1000) and incubated for 30 minutes in a dark CO2 incubator. Cells were subsequently washed with 1X PBS and immediately imaged using an EVOS imaging system (ThermoFisher Scientific). Fluorescence intensity was quantitated by using the ImageJ software^9^. The intensity of green fluorescence enables detecting cells undergoing apoptosis. Thus, cells were visualized for green fluorescence using an excitation maximum of 488 nm and, an emission maximum of 530 nm Fluorescent intensity was quantitated by using ImageJ software (National Institutes of Health, Bethesda, MA, USA). Untreated cells were used as a negative control.

### Mitochondrial Membrane Potential Detection

We used the MT1-Mito MP (Mitochondrial Membrane Potential) detection kit (Cat # MT13-10, Dojindo) to monitor changes in mitochondrial membrane potential in response to Fentanyl treatment. A549 cells were grown to ∼80% confluence in a 20mm petri dish with a glass bottom (Cat # 801001 NEST), then treated for 24 hr with Fentanyl (10-1000 ng/ml). After incubation cells were washed with 1X PBS and treated with freshly prepared MT-1 MitoMP (1:1000) and incubated for 30 minutes in a dark CO_2_ incubator. Cells were subsequently washed with 1X PBS and immediately imaged using an EVOS imaging system (ThermoFisher Scientific). Fluorescence intensity was quantitated by using the ImageJ software. The MT1-dye uptake (Detection Ex: 530-560 nm/Em: 570-640 nm) by cells was measured as red fluorescence and the intensity of dye uptake enables detecting any change in mitochondrial membrane potential. The fluorescence intensity was quantitated by using ImageJ software (National Institutes of Health, Bethesda, MA, USA). Untreated cells were used as a negative control.

### ROS quantitation

Fentanyl-induced oxidative stress was measured by quantitating the production of intracellular reactive oxygen species (ROS) in A549 treated with Fentanyl using the CM-H2DCFDA reagent from Invitrogen (Cat # C6827). CM-H2DCFDA is a chloromethyl derivative of H2DCFDA, useful as an indicator for ROS generation in cells and a general oxidative stress indicator. ROS is a green, fluorescent product that can be quantified using fluorescence microscopy imaging (Ex/Em: ∼492–495/517–527 nm). HMC-3 (10,000 cells/ml) were grown to 80% confluence in a 20mm petri dish with a glass bottom (Cat # 801001 NEST), treated with fentanyl (10-1000 ng/ml) for 24 hr, following which cells were washed with 1X PBS, following which 5μM of CM-H2DCFDA (freshly prepared in HBSS) was added, and cells were incubated for 30 min in dark CO_2_ incubator. After 30 min, cells were washed with PBS, and the ROS production was quantified by measuring the green fluorescence using the EVOS® FL Cell Imaging System (Life Technologies, Grand Island, NY). Untreated cells were used as a negative control.

### Singlet Oxygen Quantitation

A549 cells were grown to ∼80% confluence, in a 20mm petri dish with a glass bottom (Cat # 801001 NEST), then treated with Fentanyl (10-1000 ng/ml) for 24 hr. Cells were then treated with 1 µM Singlet Oxygen Sensor Green reagent (SOSG) (Cat # 22692; Lumiprobe Corporation, MD, USA) for 30 mins. Fluorescence measurements were made in a spectrofluorometer using excitation/emission wavelengths of 488/525 nm. Cells were visualized for green fluorescence, the fluorescence intensity was quantitated by using the ImageJ software (National Institutes of Health, Bethesda, MA, USA). Untreated cells were used as a negative control.

### Intracellular free Ca^2+^ concentration

A549 cells were grown to ∼80% confluence, in a 20mm petri dish with a glass bottom (Cat # 801001 NEST), then treated with Fentanyl (10-1000 ng/ml) for 24 hr. Cells were washed with HBSS buffer before being treated with 2µM Ca2+ Probe Rhod-2 AM (CAS#: 145037-81-6; AAT-Bioquest; Avantor-VWR) followed by incubation of the dye-loaded culture dish at 37 °C for 30 minutes. After incubation cells were washed with HBSS and fluorescence measurements were made in a spectrofluorometer using Ex/Em = 540/590 nm.

Cells were visualized for red fluorescence, the fluorescence intensity was quantitated by using ImageJ software (National Institutes of Health, Bethesda, MA, USA). Untreated cells were used as a negative control.

### Immunofluorescence staining of HIF-1_α_, µ opioid Receptor (MOR), and NF-_ĸ_B

A549 cells were grown to ∼80% confluence, in a 20mm petri dish with a glass bottom (Cat # 801001 NEST), then treated with Fentanyl (10-1000 ng/ml) for 24 hr, following which immunofluorescence staining was performed according to standard protocols. Briefly, cells were washed with 1X PBS, following fixation with 4% paraformaldehyde, permeabilization with 0.15% TritonX-100 (Sigma), and blocking with 3% bovine serum albumin (Sigma). Cells were stained with primary mouse monoclonal HIF1α antibody, µ opioid Receptor (MOR), and Phospho-NF-κB p65 (Ser536) (Cell signaling Technologies); and visualized using Alexa Fluor 488-conjugated or Alexa Fluor 594-conjugated anti-mouse or anti rabbit secondary antibody (Abcam) followed by staining with nuclear stain DAPI (Invitrogen). HIF1α was quantified by measuring the green fluorescence, (Alexa Fluor 488 (AF488) has an excitation peak at 488–499 nanometers (nm) and an emission peak at 519–520 nm), and MOR and NF-ĸB were quantified by measuring the red fluorescence (Alexa Fluor 594 (AF594), has an excitation peak at 590 nanometers (nm) and an emission peak at 617 nm. Using the EVOS® FL Cell Imaging System (Life Technologies, Grand Island, NY) the fluorescence intensity was quantitated by using ImageJ software (National Institutes of Health, Bethesda, MA, USA). Untreated cells were used as a negative control.

### Quantification of the relative fluorescence intensity (RFI)

Quantification of the mean fluorescence intensity (MFI) with different fentanyl concentrations was done using ImageJ software (https://imagej.net/ij/)^10^. The images were taken at three different locations for each concentration and the mean fluorescence intensity was quantified and calculated as an average of the three readings.

### Gene Expression analysis

A549 cells were treated with Fentanyl (10-1000ng/ml) concentrations for 24 hr followed by RNA extraction, cDNA synthesis and QPCR.

### RNA Extraction

RNA was extracted by an acid guanidinium-thiocyanate-phenol-chloroform method, using TRIzol® reagent. 1 ml TRIzol® reagent was added to 100,000 cells/ml A549 cells treated with Fentanyl (10-1000 ng/ml) for 24 hr. The samples were transferred to a 1.5 ml tube and 200 μl of chloroform was added. After 10 mins at room temperature, the samples were centrifuged at 13000 rpm for 15 mins at 4°C. The colorless layer was carefully transferred to a new 1.5 ml tube and 600 μl of isopropanol was added, the tubes were stored at -80°C overnight. The next day the tubes were centrifuged at 13000 rpm for 30 min at 4°C. The RNA was precipitated as a pellet and was washed twice with 1 ml of 75% ethanol followed by air drying for 5 mins and re-constitution with 25 μl of DEPC H_2_O. The amount of RNA was quantified using a Nano-Drop ND-1000 spectrophotometer and the isolated RNA was stored at -80°C.

### Real-Time Quantitative RT-PCR

According to the manufacturer’s instructions, 500 ng of the total RNA that was extracted as described above was utilized for the All-in-One Universal RT Master Mix synthesis kit (Lamba Biotech). One microliter of the resultant cDNA from the RT reaction was employed as the template in PCR reactions using well-validated PCR primers (IL1β, TNF-α, NF-κB, ERK, Caspase-3, and HIF-1α) obtained from IDT-Integrated DNA Technologies. We used the SYBR® Green master mix that contained dNTPs, MgCl_2_, and DNA polymerase (Bio-Rad, Hercules, CA). The final primer concentration used in the PCR was 0.1 µM. The following were the PCR conditions: 95°C for 3 min, followed by 40 cycles of 95°C for 40 sec, 60°C for 30 sec, and 72°C for 1 min; the final extension was at 72°C for 5 min. Gene expression was calculated using the comparative CT method. To account for variations in RNA input quantity, measurements were performed on an endogenous reference gene, β-actin. The threshold cycle (Ct) of each sample was determined, the relative level of a transcript (2ΔCt) was calculated by obtaining ΔCt (test Ct − β-actin Ct), and transcript accumulation index (TAI) was calculated (TAI = 2^-ΔΔCT^) ^11^.

### Cell Migration-Strach Assay

A549 cells were seeded in 24-well cell culture plates and cultured to a confluent monolayer. A pipette tip (10 μl) was used to scratch a wound on the midline of the culture well, and the cells were pretreated with Fentanyl (10-1000 ng/ml) for 24 hr at 37°C; the cells were then washed twice with PBS. The migration of the cells was evaluated over 4 days by measuring the number of cells migrating into the center and closing the scratched area. Imaging was done using the EVOS® FL Cell Imaging System (Life Technologies, Grand Island, NY).

### Raman Instrumentation

The spectra were measured on a DXR2 Raman microscopy setup (Thermo Fisher Scientific, Madison, WI), equipped with a laser source unit emitting ∼60 mW at 633 nm (ROUSB-633-PLR-70-1, Ondax), a 50 μm pinhole to shape the laser beam to a 0.7×0.7×1.5 μm3 FWHM, and a Plan N 100× Olympus objective lens (NA = 1.25). In addition, the Raman microscope was equipped with a fluorescence lamp (X-Cite 120 PC, Photonic Solutions) to view live cells. Fluorescence imaging was done using the EVOS® FL Cell Imaging System (Life Technologies, Grand Island, NY).

### Ramanomics

We performed Ramanomics analysis which utilizes quantitative micro-Raman spectrometry coupled with our proprietary Biomolecular Component Analysis (BCA) software to deduce composition of biomolecular constituents in subcellular domains. At first, live A549 cells were obtained in an optically transparent Dulbecco’s Modified Eagle’s Medium (DMEM) (Thermo Fisher Scientific) and mounted onto the optical stage. To ensure a high-quality signal/noise ratio, the spectra accumulation parameter was set to 6×20 s with no measurable phototoxicity after irradiation dose. During the experiments, the cells were maintained under physiological conditions at 37 °C. Spatial precision (XYZ position) of the mitochondria (30 cells/set of data) before and after each measurement was verified to attain absolute data in Raman spectra acquisition. Quantitative analysis of cellular spectra was performed using a BCAbox software (ACIS LLC, Buffalo, NY). Detailed description of the method, output (BCA) parameters and the calibration of Raman band intensities on the concentrations of biomolecules in the sample were described in our previous publications ^12,13^.

## STATISTICAL ANALYSIS

Statistical analysis was performed using GraphPad Prism (v8; GraphPad). Differences between fentanyl-treated and control samples were assessed using a two-tailed Student t-test. For multiple pairwise comparisons, data were analyzed using one-way ANOVA, followed by Tukey’s multiple comparison test. Statistical significance was defined as P < 0.05.

## RESULTS

### Effect of Fentanyl on Cell Viability in A549 cells

Cell viability was determined using CCK-8 (Cell Counting Kit-8) assay following the the manufacturer’s protocol. The observation (Fig 1a) shows that, Fentanyl at doses from 1-100 ng/ml had a cell viability > 95% and a significant decrease to 75% at the fentanyl concentration of 1000 ng/ml (p=0.0001).

**Figure 1.**
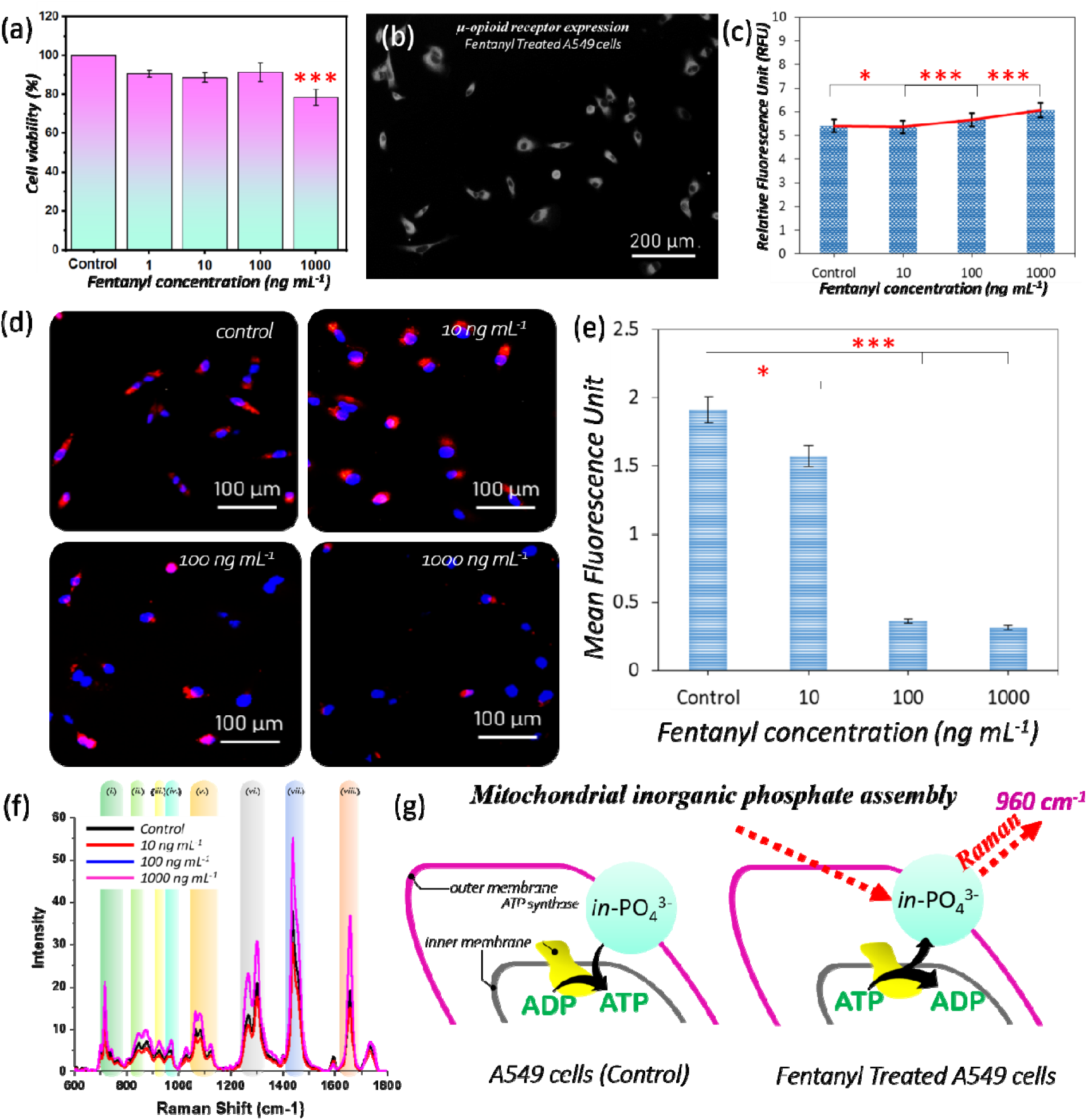
Cellular and mitochondrial responses of A549 cells to fentanyl exposure. (a) MTT assay measuring A549 cell viability following treatment with increasing concentrations of fentanyl. Statistical significance indicates ***p<0.0001 between untreated control and 1000ng/ml concentration fentanyl treatment. (b) Representative imag of Immunofluorescence detection of μ-opioid receptor expression in A549 cells. (c) Relative fluorescence intensity of μ-opioid receptor expression across varying fentanyl doses as compared to the control. Statistical significance indicates *p<0.05 and ***p<0.0001; (d) Representative fluorescence images of mitochondrial staining in A549 cells treated with increasing concentrations of fentanyl. (e) Quantitative analysis of mitochondrial membrane potential based on fluorescence intensity across fentanyl treatments with respect to the untread control; Statistical significance indicates *p<0.05 and ***p<0.0001 (f) Pre-processed lipid profile of mitochondria extracted from A549 cells. (g) Schematic overview depicting the inorganic phosphate signature observed in the Ramanomics analysis on comparison between untreated control and fentanyl treated A549 cells highlighting its association with mitochondrial membrane potential.

### Immunofluorescence staining of µ opioid receptor in A549 cells

Treatment of A549 cells with fentanyl elicited a concentration-dependent increase in μ-opioid receptor fluorescence (λ_ex = 488 nm; Fig. 1b). Quantification of mean fluorescence intensity (MFI) using ImageJ (NIH, Bethesda, MD, USA) revealed a 35-fold increase at 10 ng/mL (p < 0.05), a 65-fold increase at 100 ng/mL (p < 0.001), and a 55-fold increase at 1000 ng/mL (p < 0.001) compared to untreated controls (Fig. 1c). Statistical significance was defined as p < 0.05.

### Effect of Fentanyl on Mitochondrial Membrane Potential

To assess the impact of fentanyl on mitochondrial membrane potential (Δψm), A549 cells were stained with MT1-Mito MP and imaged (λ_ex/λ_em 488/610 nm). Quantitative analysis of mean fluorescence intensity (MFI) revealed that cells exposed to 100 and 1000 ng·mL□^1^ fentanyl exhibited an ∼85% reduction in Δψm relative to untreated controls (Fig. 1d; p < 0.001). The profound collapse of Δψm indicates dissipation of the proton motive force that drives ATP synthase. In support of this, Raman spectral mapping of regions of interest (Fig. 1g) and corresponding proton/inorganic phosphate vibrational bands (Fig. 1g) revealed impaired ADP⇌ATP interconversion in fentanyl-treated cells. These findings link fentanyl-induced Δψm loss to bioenergetic failure and altered mitochondrial dynamics changes that can compromise ATP production, enhance reactive oxygen species generation, and sensitize cells to apoptotic signaling (Fig. 1e).

### Raman spectra of mitochondrial fractions from fentanyl-treated A549 cells

Pre-processed, baseline□corrected Raman spectra of mitochondrial fractions from fentanyl-treated A549 cells revealed dose-dependent shifts in bands assigned to nucleic acids (Fig. 1f i, vi), proteins/carbohydrates (Fig. 1f ii–v) and lipids (Fig. 1f vii–viii). Because fentanyl is highly lipophilic and accumulates within membrane bilayers, we focused on the lipid associated spectral region to interrogate changes in the outer and inner mitochondrial membranes. Analysis of lipid vibrational peaks—particularly those reflecting acyl□chain order (∼1,300–1,450 cm□^1^) and phospholipid headgroup composition (∼720–800 cm□^1^)—uncovered marked alterations in membrane chemical configuration with increasing fentanyl dose. These findings are relevant because re-modeling of mitochondrial membrane lipids can modulate membrane fluidity, permeability and the function of embedded respiratory complexes, linking fentanyl’s lipophilicity to its effects on mitochondrial bioenergetics and downstream apoptotic signalling (Fig. 1g).

### Effect of Fentanyl on NF-**ĸ**B in A549 cells

We evaluated the impact of fentanyl on NF-κB signaling in A549 cells by measuring p65 phosphorylation, and NF-κB–driven transcription. Immunofluorescence staining of Phospho-NF-κB p65 was done in fentanyl-treated A549 cell line at dilution of 1:1000 and secondary antibody staining using Alexa Fluor 594 Anti-Rabbit IgG at 1:5000 dilution. Fig. 2a-d show an increase in Phospho-NF-κB p65 expression confirmed rapid nuclear translocation of p65 after fentanyl exposure as seen with increase in red fluorescence emission with increase in fentanyl dose.

**Figure 2.**
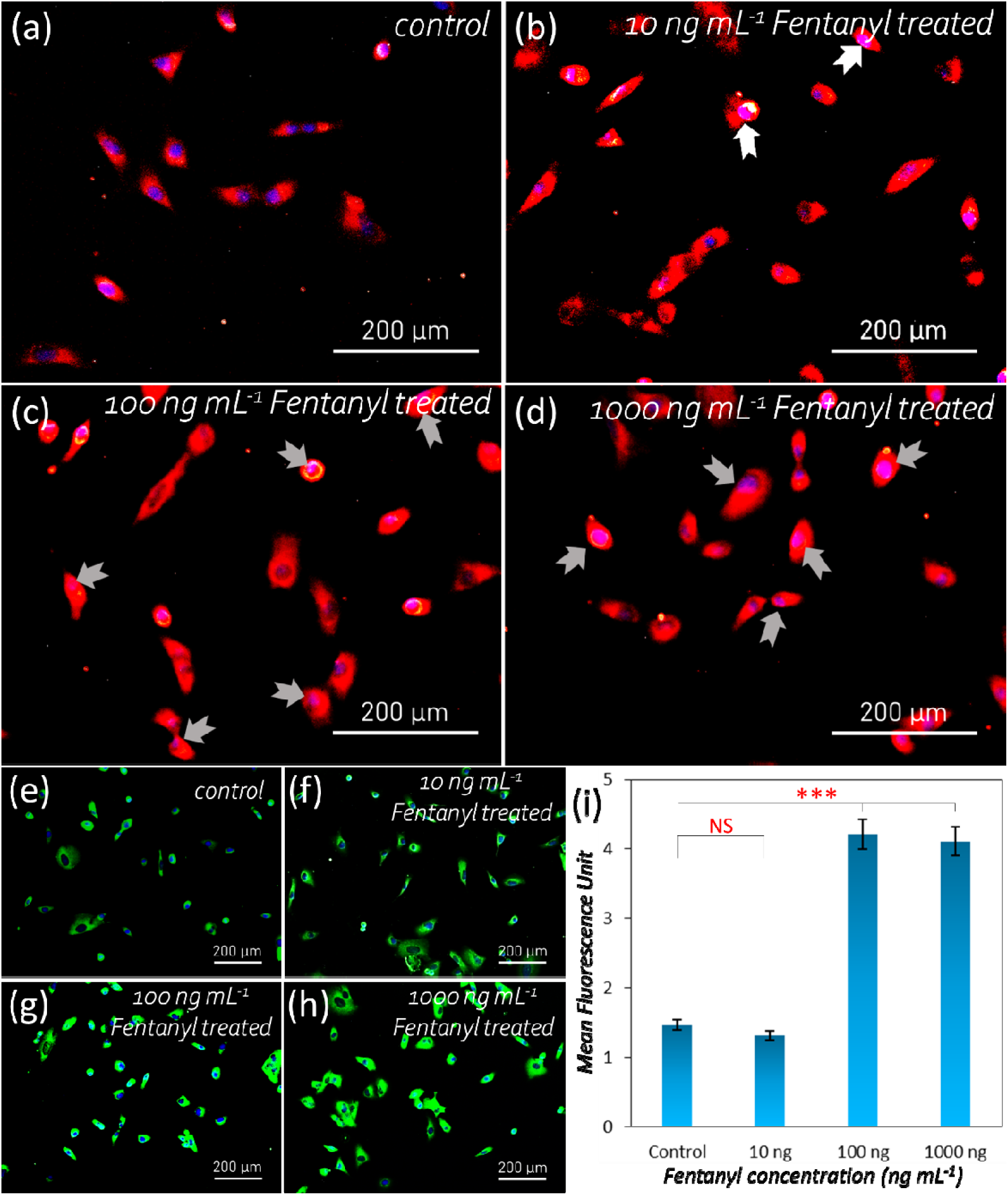
Dose-dependent activation of inflammatory and hypoxic pathways in A549 cells following fentanyl exposure. (a–d) Representative Immunofluorescence images showing nuclear translocation and activation of NFκB in A549 cells treated with increasing concentrations of fentanyl.(e, h) Representative fluorescence images displayin HIF-1α expression in A549 cells treated with 10, 100, and 1000 ng/mL fentanyl, compared to untreated controls.(i) Quantitative plot of relative fluorescence intensity illustrating hypoxia generation across varying fentanyl dosages. Statistical significance indicates *p<0.0001; NS (not significant)

### Immunofluorescence staining of HIF-1α in A549 cells

Our data shows a dose-dependent increase in HIF-1α expression in response to Fentanyl treatment in A549 cells (Fig. 2e-h). We observed an increase in the green fluorescence intensity at higher fentanyl concentrations compared to the untreated control. We observed a 69% (p<0.0001) and 63% (p<0.0001) increase in HIF-1α expression in A549 cells treated at 100 and 1000 ng/mL fentanyl concentrations and no significant differences in HIF-1α expression at 10 ng/ml Fentanyl concentrations as compared to the untreated control (Fig. 2i) .

### Effect of Fentanyl on Reactive Oxygen Species (ROS) production

Intracellular ROS levels were quantified using the DCFH-DA probe by fluorescence microscopy. A549 cells treated for 24 h with fentanyl at 10, 100 and 1000 ng·m□^1^ exhibited a dose-dependent increase in ROS signal.

Representative fluorescence micrographs of DCFH-DA–stained A549 cells under control conditions (Fig. 3a-d) and following treatment with 1000 ng/mL fentanyl reveal a marked increase in intracellular ROS. Quantitative analysis of mean fluorescence intensity (MFI), performed in ImageJ, is presented in Fig. 3e, We observed a 32% (p<0.05) and 27% (p<0.05) increase in ROS levels in 100 and 1000 ng/ml fentanyl concentrations as compared with untreated controls, while no significant difference was observed when A549 cells were treated with 10ng/ml fentanyl when compared to the untreated control. These data demonstrate that fentanyl markedly elevates oxidative stress in A549 cells, a change that may underlie its effects on mitochondrial function, HIF-1α stabilization and activation of redox-sensitive signaling pathways.

**Figure 3.**
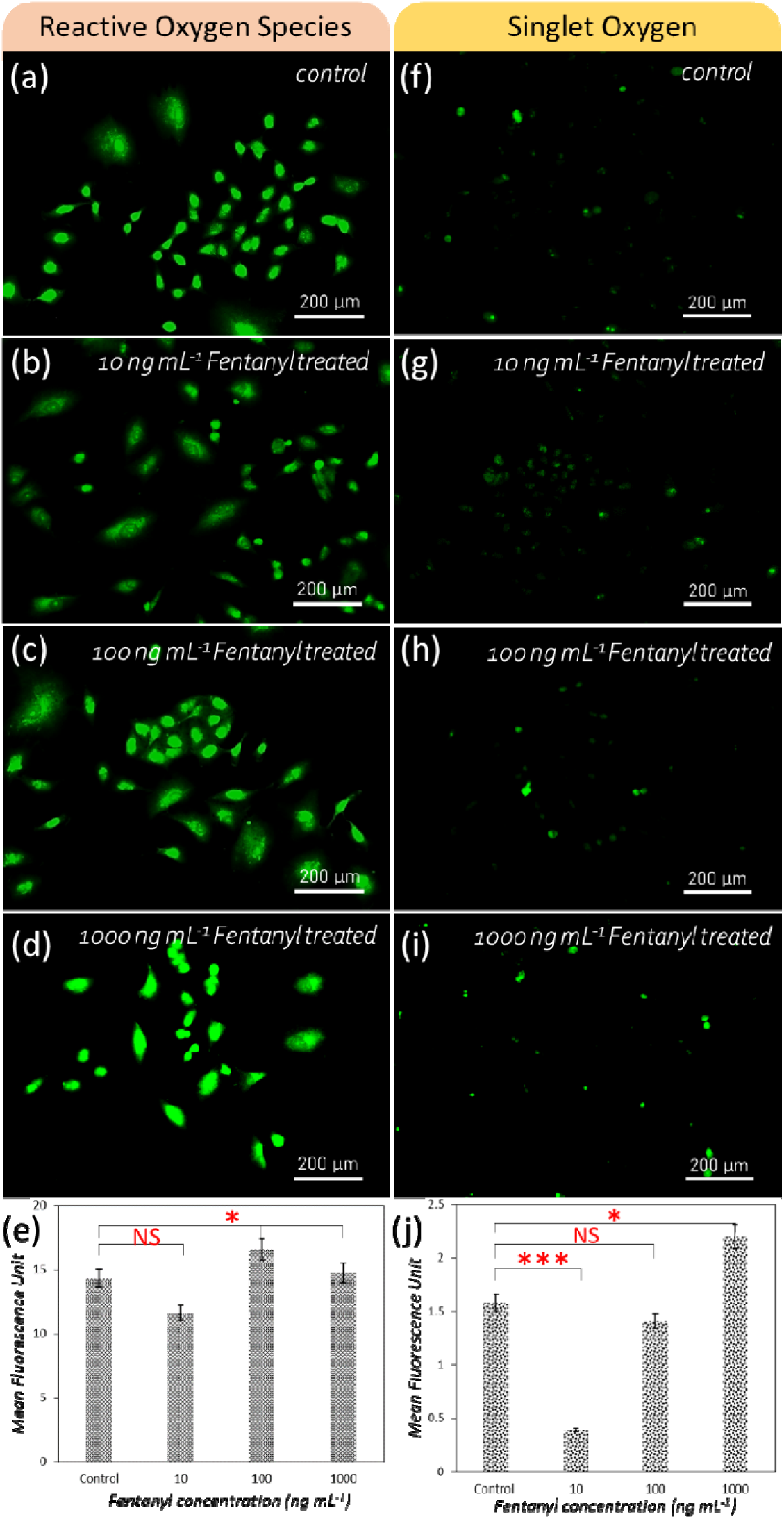
Fentanyl-induced oxidative stress in A549 cells. (a–d) Representative fluorescence images showing reactive oxygen species (ROS) accumulation in A549 cells following treatment with increasing concentrations of fentanyl.(e) Quantitative plot of relative fluorescence intensity for ROS across various fentanyl doses. . Statistical significance indicates **p<0.001(f–i) Representative fluorescence images displaying singlet oxygen species in A549 cells treated with increasing fentanyl concentrations. (j) Relative fluorescence plot quantifying singlet oxygen generation at corresponding fentanyl dosages with respect to the untreated control. Statistical significance indicates *p<0.05 and ***p<0.0001; NS (not significant)

### Effect of Fentanyl on Singlet Oxygen production

Singlet oxygen production was assessed in A549 cells using the Singlet Oxygen Sensor Green (SOSG) probe and fluorescence microscopy. Treatment with 1000 ng·mL□^1^ fentanyl for 24 h elicited a 37.5 % increase (p < 0.05), in singlet oxygen signal relative to untreated controls whereas treatment of A549 cells with 10 ng·/mL fentanyl reduced singlet oxygen levels by approximately 70 % (p < 0.0001) whereas no significant changes in Singlet oxygen production was observed at fentanyl treatment of 100ng/ml when compared to the untreated control. (Fig. 3f–i). These changes in Singlet oxygen production indicate that fentanyl modulates not only bulk ROS but also the specific reactive oxygen species profile, with potential implications for oxidative damage and redox□sensitive signaling in lung cancer cells. These data suggest that fentanyl could be influencing enzymes or mediators that modulate singlet oxygen levels, and those pathways might be more sensitive at low doses and higher fentanyl decrease more effectively.

### Effect of Fentanyl on intracellular Ca^2+^

Intracellular Ca2+ was measured in Fluo-4 AM– loaded A549 cells by fluorescence microscopy. Representative red-fluorescence images for control and fentanyl-treated (10, 100, 1000 ng/mL) cells are shown in Fig. 4a–d. Quantification of mean fluorescence intensity demonstrated that 10 ng/mL fentanyl reduced intracellular Ca2+ by 72% (p < 0.001), versus untreated cells whereas higher doses of Fentanyl produced increase in Ca2+, of 33% (p < 0.05) and 36%(p < 0.05) increase at a fentanyl concentration of 100 and 1000 ng/mL respectively as compared to the untreated control (Fig. 4e).

**Figure 4.**
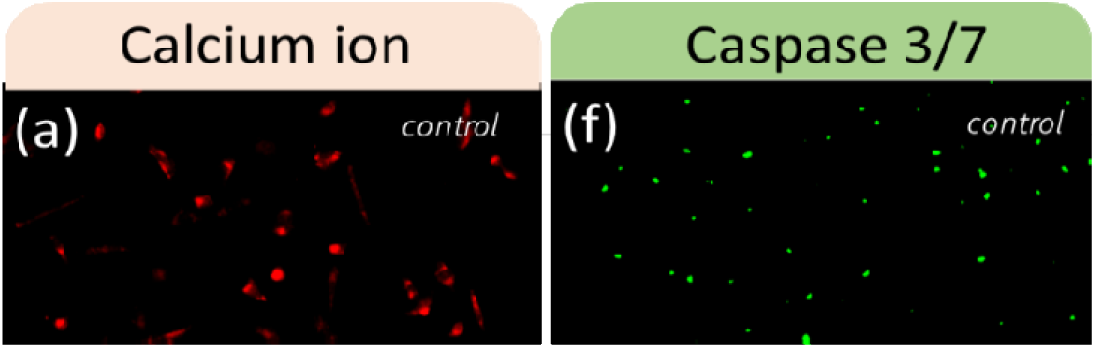
Modulation of calcium signaling and apoptotic enzyme activity in A549 cells following fentanyl exposure. (a–d) Representative Fluorescence images illustrating calcium ion distribution in A549 cells treated with increasing doses of fentanyl. (e) Quantitative plot of relative fluorescence intensity depicting changes in calcium ion signaling across fentanyl concentrations with respect to the untreated control. Statistical significance indicates *p<0.05 (f–i) Representative fluorescence images showing Caspase 3/7 enzyme activation in A549 cells in response to increasing doses of fentanyl (j) Relative fluorescence intensity plot quantifying Caspase 3/7 levels at corresponding fentanyl doses with respect to the untreated control. Statistical significance indicates *p<0.05; **p<0.001 and ***p<0.000; NS (not significant)

### Effect of Fentanyl on Caspase 3/7 Activity

Caspase-3/7 activation in A549 cells was assessed by fluorescence staining of a fluorogenic caspase substrate (Fig. 4f–i). We observed no significant change in Caspase 3/7 activation at a Fentanyl dose of 10ng/ml, however, we observed a 68% ((p < 0.0001) and 72% ((p < 0.0001) increase in caspase-3/7 activity when A549 cells were treated with a fentanyl dose of 100 ng/ml and 100 ng/ml respectively as compared to the untreated control (Fig. 4j).. Our data indicate that fentanyl at elevated concentrations robustly activates the executioner caspases, promoting apoptotic signaling in lung cancer cells.

### Effect of Fentanyl on Gene Expression of Pro-inflammatory Cytokines, Hypoxia-Related, and Pro-Apoptotic Transcriptional Regulators

Quantitative RT-PCR was used to investigate the impact of fentanyl on the expression of selected genes associated with inflammation, hypoxia response, apoptosis, and signaling in A549 cells. The analysis included IL-1β, TNF-α, NF-κB, Caspase-3, HIF-1α, and ERK1/2, with gene expression quantified using the comparative CT (ΔΔCT) method and reported as transcript accumulation index (TAI = 2^−ΔΔCT). Data represent the mean ± SD from three independent experiments, with statistical significance set at p < 0.05.

### Fentanyl treatment elicited dose-dependent changes in gene expression

**IL-1**β gene expression was 3.9 fold (p<0.001), 3.0 fold (p<0.001) and 5.13 fold (p < 0.0001) higher at 10, 100 and 1000ng/ml fentanyl concentrations as compared to the untreated control indicating a robust pro-inflammatory response (Fig 5a).

**Figure 5.**
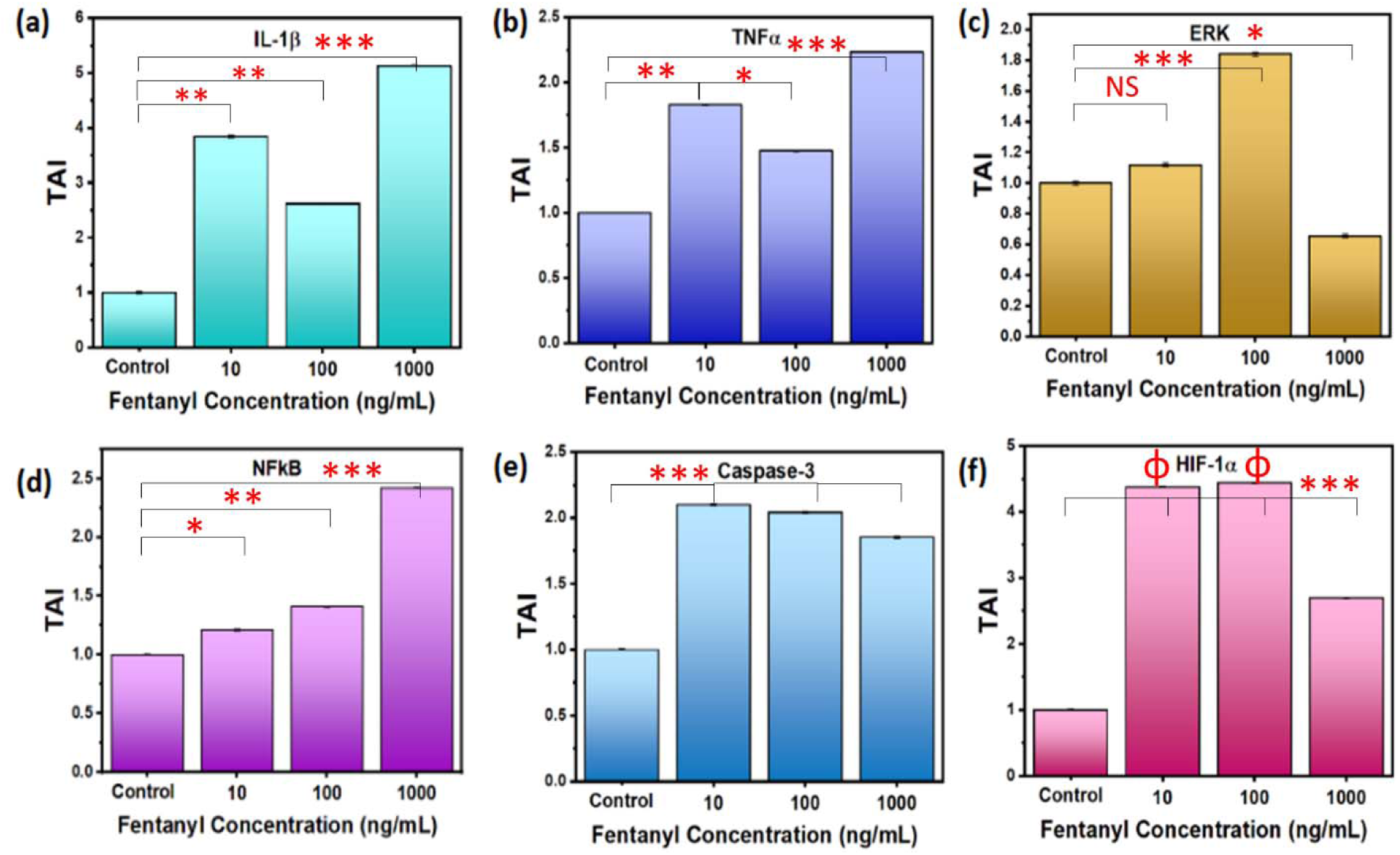
Effect of fentanyl on inflammatory, apoptotic, and hypoxic gene expression in A549 cells. (a–f) Quantitative analysis of gene expression levels for IL-1β (a), TNF-α (b), ERK (c), NFκB (d), Caspase 3 (e), an HIF-1α (f) in A549 cells treated with increasing concentrations of fentanyl. Gene expression was measured using the comparative Ct (ΔΔCt) method. The threshold cycle (Ct) of each target gene was normalized to β-actin (ΔCt), an the transcript accumulation index (TAI) was calculated as TAI = 2^−ΔΔCt. Data are presented as mean ± SD from three independent PCR experiments (n = 3). Statistical significance indicates *p<0.05, **p<0.001, ***p<0.0001, ΦP<0.000001; NS (not significant)

**TNF-**α gene expression was 1.83-fold (p < 0.001), 1.47-fold (p < 0.05), and 2.23-fold (p < 0.0001) at 10, 100 and 1000ng/ml fentanyl concentrations as compared to the untreated control further supporting a pro-inflammatory effect. (Fig 5b).

**ERK1/2** gene expression showed a biphasic response not showing any significant increase at 10 ng/mL (p=NS), but showing a 84 % increase at 100 ng/mL fentanyl (p < 0.0001), and notable downregulation at a fentanyl dose of 1000 ng/mL (34%, p< 0.05) with respect to the untreated control. (Fig 5c).

**NF-**κ**B** gene expression showed a dose dependent increase 29% (p < 0.05), 41% (p < 0.001), and 142% (p <0.0001) at 10, 100 and 1000ng/ml fentanyl concentrations respectively as compared to the untreated control consistent with inflammatory signaling. Fig 5d).

**Caspase-3** gene expression showed a dose dependent increase 110% (p < 0.0001), 104%(p < 0.0001), and 85% (p < 0.0001) at 10, 100 and 1000ng/ml fentanyl concentrations respectively as compared to the untreated control highlighting activation of apoptotic mechanisms. Fig 5e).

**HIF-1**α gene expression showed significant increase of 338% (p < 0.000001), 343% (p < 0.000001) and 169% (p < 0.0001) at 10, 100 and 1000ng/ml fentanyl concentrations respectively as compared to the untreated control indicating significant activation of hypoxia-associated pathways. Fig 5f).

These findings demonstrate that fentanyl, particularly at higher doses, induces a transcriptional signature consistent with pro-inflammatory, hypoxia-responsive, and pro-apoptotic cellular stress in A549 cells. Fentanyl’s dose-dependent activation of pro-inflammatory and pro-apoptotic gene pathways (like IL-1β, TNF-α, NF-κB, and Caspase-3), along with elevated hypoxia signaling (HIF-1α), suggests that it induces cellular stress and may compromise cell survival. These effects indicate that fentanyl actively reprograms cellular behavior. The biphasic response of ERK1/2 hints at a complex signaling dynamic, possibly tied to feedback mechanisms or differing thresholds for cell adaptation versus stress-induced shutdown.

### Effect of Fentanyl on A549 Cell Migration

Wound-healing (scratch) assays were conducted to assess the effect of fentanyl on A549 cell motility. A uniform gap was created in confluent cell monolayers, followed by treatment with fentanyl at concentrations of 0 (control), 10, 100, and 1,000 ng/mL. Phase-contrast images were captured daily from Day 1 through Day 4 (Fig. 6), and gap closure was quantitatively analyzed. Fentanyl impaired cell migration in a dose-dependent manner. At 1,000 ng/mL, the wound closure was significantly reduced compared to untreated controls, showing decreases of 27% (p<0.05), 34% (p<0.05), 31% (p<0.05), and 21% (p<0.05) on Days 1, 2, 3, and 4, respectively. By Day 4, control cells had nearly fully closed the wound, while fentanyl-treated cultures demonstrated on an average approximately 25% less closure. (Fig. 6b). Fentanyl reduces cell motility in a concentration-dependent manner. The higher the fentanyl dose, the more impaired the migration, with the most dramatic effects seen at 1,000 ng/mL. At the fentanyl dose of 1,000 ng/mL, wound closure lagged consistently behind the control over all four days. Even by Day 4, cells treated with fentanyl showed ∼25% less gap closure than untreated cells, suggesting persistent suppression of migratory capacity. Data are presented as mean ± SD from three independent experiments; p < 0.05 was considered statistically significant.

**Figure 6.**
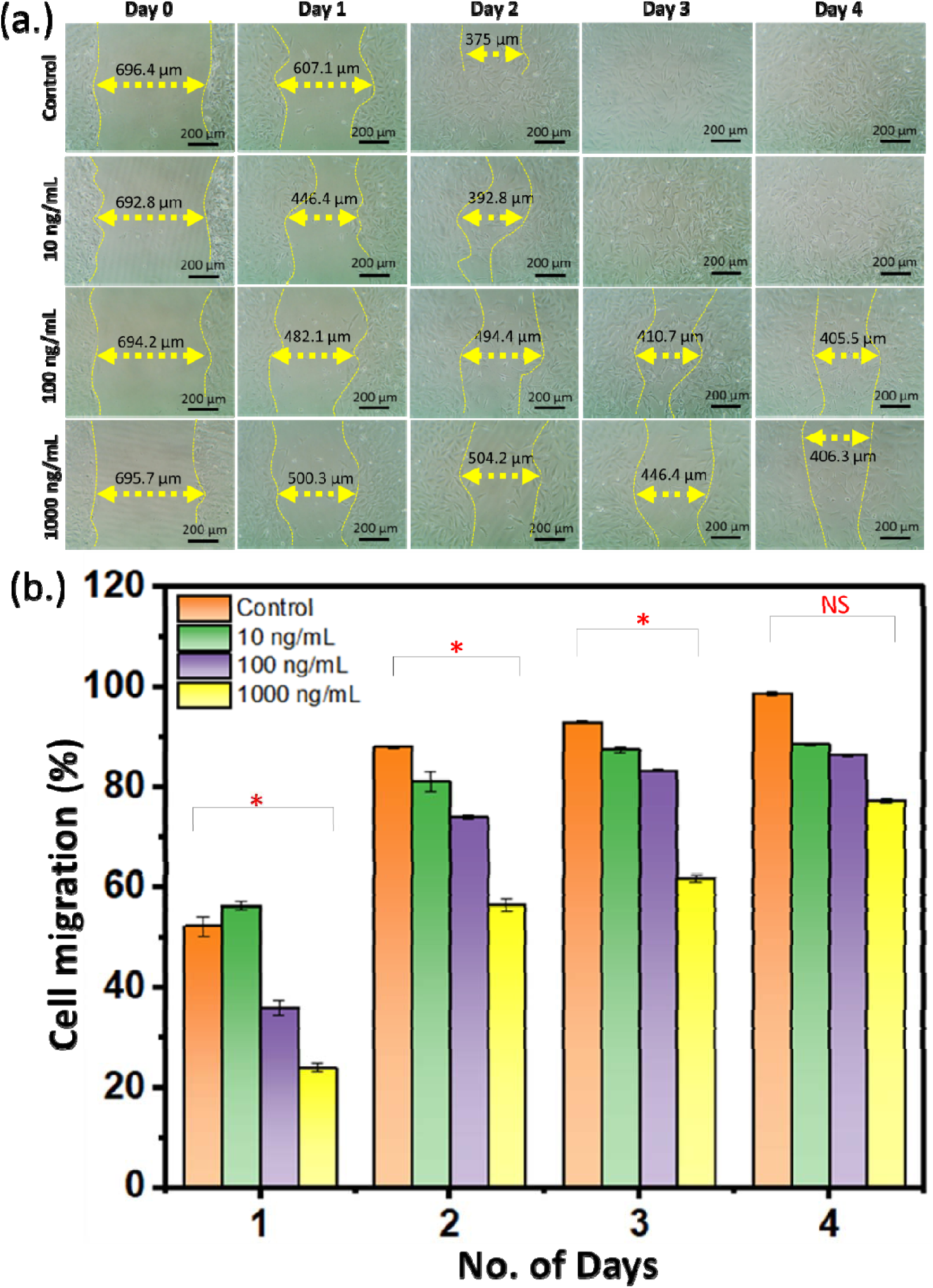
Impact of fentanyl on cell migration and proliferation in A549 cells. (a) Bright-field microscopy images depicting cell migration patterns in A549 cells treated with various concentrations of fentanyl compared t untreated controls. (b) Comparative analysis of cell proliferation behavior in A549 cells following drug treatment, presented as a quantitative plot. Statistical significance indicates *p<0.05; NS (not significant)

### Ramanomics study for fentanyl overdose in A549 cells

Figure 7a presents a schematic illustration of the Ramanomics assay, derived from stained mitochondrial analysis in A549 cells. The study examined changes in carbohydrate (glycogen), sterols (cholesterol, ergosterol, cholesterol esters), lipids (phospholipids), nucleic acids (DNA, t-RNA, m-RNA), total protein, and phosphocholine in response to fentanyl exposure.

**Figure 7.**
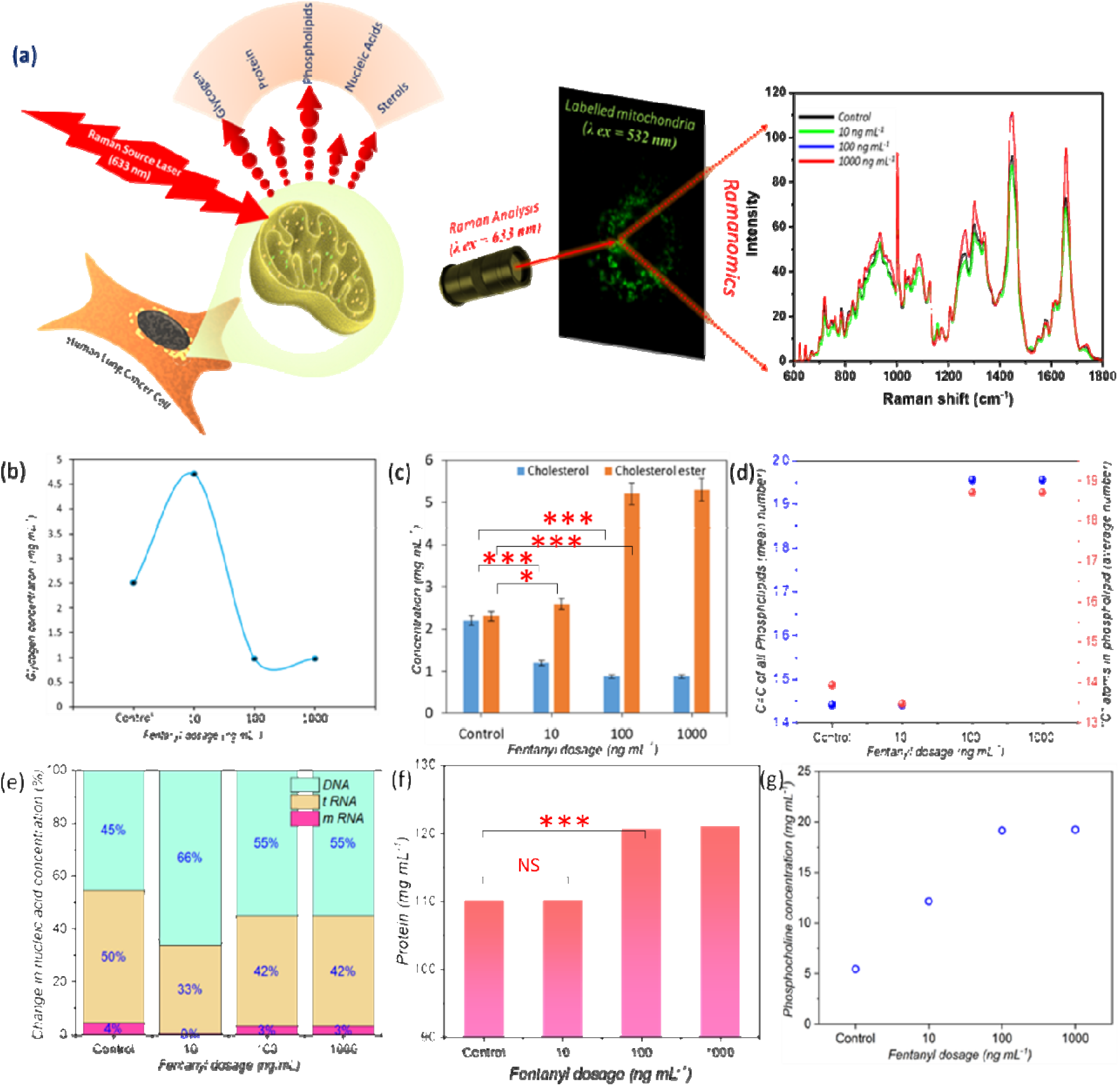
Raman-based biochemical profiling of mitochondria in A549 cells following fentanyl exposure. (a) Schematic representation of the Raman spectroscopy workflow used for mitochondrial analysis in fentanyl-treated A549 cells. (b–g) Quantitative Ramanomic assessment of biochemical changes in mitochondria corresponding to increasing fentanyl concentrations:(b) glycogen content;(c) sterol levels;(d) saturation status of phospholipids;(e) mitochondrial nucleic acid composition;(f) mitochondrial protein content; (g) phosphocholine concentration. Statistical significance indicates *p<0.05, *** p<0.0001; NS (not significant)

Glycogen levels initially increased (∼1.8-fold) but subsequently declined (∼2.5-fold) upon treatment with 100–1000 ng/mL of fentanyl (Fig. 7b). Within the sterol category (Fig. 7c), cholesterol and ergosterol concentrations decreased by 2.7-fold and 2.1-fold, respectively, in response to fentanyl overdose. Conversely, cholesterol ester levels exhibited a 2.5-fold increase under the same dosage range.

Phospholipid analysis revealed an increase in the mean C=C ratio of phospholipids from ∼1.5 to 2, along with a rise in the total carbon content, indicating enhanced unsaturation within the mitochondrial environment (Fig. 7d). Nucleic acid content, mitochondrial DNA levels increased by 10–20% following fentanyl exposure, whereas t-RNA levels showed an average 12.5% decrease (Fig. 7e).

Total mitochondrial protein content increased from 109.5 mg/mL to 120 mg/mL following fentanyl overdose, while a lower dose (10 ng/mL) produced no significant change. Additionally, phosphocholine, a signaling molecule, exhibited a gradual increase from ∼5 mg/mL to 20 mg/mL, reflecting a 4-fold elevation in mitochondrial phosphocholine levels in A549 cells.

## DISCUSSION

Fentanyl induced hypoxia and oxidative stress are closely linked to metabolic reprogramming in cancer cells and these conditions can alter pathways like glycolysis and oxidative phosphorylation, potentially affecting the growth rate and survival of lung cancer cells. The goal of this study was to explore the effects of fentanyl on ROS, singlet oxygen generation, mitochondrial activation, expression of hypoxia inducing factors, activation of apoptosis, and cytokine production in A549 cells and examine the potential mechanisms that underlie the fentanyl-induced effects. The findings associated to fentanyl overdose in human lung cancer (A549) cell line reveal biochemical changes at organelle and molecular level.

Our MTT assay shows a 25% decrease in cell viability upon exposure to the highest fentanyl dose of 1000ng/ml and no significant cell toxicity at lower doses of fentanyl (10-100 ng/ml). All experimental paradigms were conducted at the Fentanyl dose range of 10-1000 ng/ml.

Immunofluorescence staining of A549 cells with μ-opioid receptor antibody showed an increased expression of μ-opioid receptor in fentanyl treated A549 cells. This confirms the presence of μ-opioid receptor on the cancer cell line which promotes fast uptake of fentanyl resulting in activation of signaling pathways that promote cancer cell migration and cell proliferation in human lung tumor-associated endothelial cells as observed by Liu et al ^14^.

Mitochondrial membrane potential staining provides crucial insights into mitochondrial function, revealing a decrease in treated cells compared to untreated controls. We showed an ∼85% reduction in mitochondrial membrane potential (Δψm) at high fentanyl doses, along with mitochondrial fusion and impaired ADP/ATP conversion. This reinforces that fentanyl impairs mitochondrial bioenergetics and cell membrane integrity. Correlating these findings with the mitochondrial lipid model derived from the pre-processed dataset, a notable shift in the Raman peak at 960 cm□^1^ (Fig. 1f), corresponding to inorganic phosphate (in-PO□^3^□), was observed across varying fentanyl concentrations. This suggests that the conversion of ATP to ADP was reversed (ADP to ATP) at a fentanyl concentration of 10 ng/mL, whereas excessive fentanyl exposure (100–1000 ng/mL) further promoted this effect. The inorganic phosphate (in-PO□^3^□) released during ATP hydrolysis accumulated within the inner mitochondrial membrane, contributing to mitochondrial dysfunction and a significant reduction in Δψm (Fig. 1e) in cells exposed to higher fentanyl concentrations. Our study highlights potential pro-tumorigenic risks via mitochondrial destabilization and oxidative stress, suggesting that fentanyl’s effects may be context- and dose-dependent.

Immunofluorescent staining of the NF-κB p65 subunit in A549 cells revealed a dose-dependent increase in both total and nuclear fluorescence intensity following fentanyl exposure (Fig. 2a-d). This upregulation of NF-κB coincides with enhanced transcription of pro-inflammatory cytokines (IL-1β, TNF-α) and activation of apoptotic effectors, underscoring NF-κB’s central role in linking fentanyl-induced redox stress to inflammatory gene expression and programmed cell death.

Our study demonstrates that fentanyl treatment in A549 cells resulted in an ∼ 67% increase in HIF-1α expression. This upregulation may contribute to elevated reactive oxygen species (ROS) levels by modulating mitochondrial activity and enhancing the expression of enzymes such as NADPH oxidase, which is responsible for ROS generation^15^. We observed elevated ROS and singlet oxygen levels in fentanyl-treated A549 cells, contributing to redox imbalance and potential DNA damage. Fentanyl induces oxidative stress, which may contribute to cytotoxicity and altered signaling in lung cancer cells. Fentanyl-induced apoptosis in A549 cells, likely mediated by mitochondrial dysfunction and ROS accumulation. Fentanyl induces ROS production in A549 cells. It was reported that fentanyl induces ROS which contributes to autophagy in A549 cells^17^. ROS is generated by way of normal oxidative metabolism and plays a role in cell signaling and homeostasis, however, increased levels of ROS cause irreversible oxidative damage to cellular structures^18^. Fentanyl can induce the production of ROS in A549 lung cells. This increase in ROS can lead to oxidative stress, which can damage cellular components such as DNA, proteins, and lipids. Fentanyl induced increase in ROS levels can affect cell viability and promote apoptosis or programmed cell death^19^.

ROS levels in cancer cells are typically elevated due to increased cellular metabolism; however, our study specifically evaluated the contribution of singlet oxygen (^1^O□). At an optimal fentanyl concentration of 10 ng/mL, ^1^O levels decreased by 70%, while its relative contribution to the overall ROS pool increased to 37.5%. Ramanomics analysis (Fig. 7) revealed that ^1^O□ actively degraded glycogen, a primary energy source for cancer cell proliferation, in close proximity to mitochondria. Excessive singlet oxygen generation contributes to oxidative stress and cellular toxicity in A549 cells, exacerbating fentanyl-induced mitochondrial dysfunction.

The mechanisms underlying hypoxia-driven singlet oxygen production involve electron transport chain (ETC) disruptions and mitochondrial impairment.^16^ Under hypoxic conditions, inefficient ETC function leads to electron accumulation in mitochondrial complexes, facilitating the partial reduction of oxygen and subsequent ROS formation, including ^1^O□. Hypoxia-inducible factors (HIFs) further modulate mitochondrial metabolism, promoting glycolysis while reducing oxidative phosphorylation. This metabolic shift increases electron leakage, exacerbating ROS production. Additionally, impaired oxygen availability compromises cytochrome c oxidase (Complex IV), amplifying mitochondrial stress and oxidative damage. Hypoxic conditions may stabilize HIFs in lung cancer cells, thereby altering metabolic pathways, intensifying oxidative stress, and potentially promoting tumor progression and therapy resistance. HIF-1α drives angiogenesis, glycolytic reprogramming and chemoresistance in tumors, its μ-opioid–mediated activation by fentanyl may promote pseudo-hypoxic signaling and metabolic shifts that influence lung cancer progression and therapeutic response.

Elevated intracellular Ca2+ has far-reaching consequences for cancer cell physiology. As a versatile second messenger, Ca2+ modulates mitochondrial function by sensitizing the permeability transition pore, driving Δψm collapse and enhancing ROS generation. It also activates Ca2+-dependent kinases (e.g., CaMKII), phosphatases (e.g., calcineurin), and proteases (e.g., calpains), which converge on transcription factors such as NF-κB and CREB to regulate genes controlling proliferation, survival and inflammation. Thus, the fentanyl-induced Ca2+ surge at high doses likely underpins the mitochondrial depolarization, oxidative stress and apoptotic signaling we observed. Conversely, the Ca2+ reduction seen at low fentanyl concentrations may transiently dampen these pathways, illustrating a biphasic, dose-dependent regulation of lung cancer cell fate by μ-opioid receptor signaling.

A dose-dependent increase in Ca^2^□ response was observed (Fig. 4e), indicating that fentanyl (100–1000 ng/mL) upregulates intracellular Ca^2^□ by promoting Ca^2^□ influx in A549 cells. Evidence suggests that this fentanyl-induced Ca^2^□ response is mediated through mu opioid receptors expressed on A549 cells, which play a pivotal role in modulating cellular responses to fentanyl. Upon fentanyl binding, the mu receptor initiates a signaling cascade that leads to the release of Ca^2^□ from intracellular stores, a process essential for various cellular functions, including survival, proliferation, and apoptosis. ^20,21^. Fentanyl can inhibit voltage-gated calcium channels, reducing neurotransmitter release, however in cancer cells, fentanyl-induced calcium signaling may contribute to apoptosis or altered cellular metabolism^20,21^.

Fentanyl-induced cell death was found to be dependent on Caspase-3 and Caspase-7, key mediators of the apoptotic pathway. Our gene expression analysis (Fig. 5) revealed significant upregulation of Caspase-3 expression in response to fentanyl treatment at doses ranging from 100–1000 ng/mL. As a primary executioner caspase, elevated Caspase-3 expression indicates active apoptotic processes, suggesting that fentanyl treatment induces programmed cell death. Caspase-7, functioning in a complementary role, reinforces apoptosis by targeting structural and regulatory proteins. Studies indicate that fentanyl exposure can initiate apoptosis via mitochondrial dysfunction and oxidative stress, leading to Caspase-3/7 activation. This mechanism is particularly relevant in fentanyl-induced neurotoxicity and its potential impact on cancer cells. Furthermore, caspase-mediated apoptosis plays a critical role in modulating drug resistance and sensitivity in cancer therapy ^22^.

Our study observed a significant increase in Hypoxia-Inducible Factor 1-alpha (HIF-1α) gene expression, a key regulator of cellular responses to hypoxic conditions. The data indicate that fentanyl treatment induces hypoxia, subsequently triggering the upregulation of HIF-1α. This transcription factor plays a crucial role in cellular adaptation to oxygen deficiency by modulating genes involved in metabolism, angiogenesis, and survival. Under hypoxic conditions, mitochondrial dysfunction and electron transport chain (ETC) disruptions contribute to elevated reactive oxygen species (ROS) production, further stabilizing HIF-1α. This stabilization enhances its transcriptional activity, potentially influencing metabolic reprogramming and cellular stress responses in fentanyl-treated A549 cells ^23^.

Fentanyl induced respiratory depression may create localized hypoxia, stabilizing HIF-1α and altering transcription of genes that govern cytoskeletal dynamics, extracellular matrix remodeling and angiogenesis. These pseudo-hypoxic conditions likely underlie the observed reduction in A549 cell migration that we observed in our cell migration assay (Fig 6). This suggests that fentanyl may interfere with cytoskeletal dynamics or signaling pathways critical for migration.

Fentanyl treatment resulted in a significant upregulation of pro-inflammatory cytokines, including IL-1β, TNF-α, and transcription factor NF-κB, indicating that fentanyl induces a robust pro-inflammatory response in A549 cells at concentrations ranging from 100–1000 ng/mL. IL-1β and TNF-α, critical regulators of inflammation and immune signaling, exhibited increased gene expression levels following fentanyl exposure, leading to elevated cytokine production and subsequent activation of key signaling pathways, including ERK and NF-κB. These pathways play a vital role in modulating cellular responses to fentanyl. Further a dose-dependent, biphasic regulation of ERK suggests that moderate fentanyl exposure transiently engages the ERK arm of the MAPK cascade, potentially promoting pro-survival or stress-adaptive signaling, whereas high concentrations attenuate ERK activity, contributing to cytotoxic stress. Fentanyl has been shown to influence ERK (extracellular signal-regulated kinase) signaling in A549 lung cells. ERK signaling, an integral component of the mitogen-activated protein kinase (MAPK) pathway, is crucial for regulating cell proliferation, differentiation, and survival. The modulation of ERK signaling by fentanyl is predominantly mediated through its interaction with mu opioid receptors (MORs) expressed on A549 cells. Upon binding, fentanyl initiates a cascade of intracellular events, culminating in the activation of ERK1/2. The fentanyl-induced activation of ERK signaling may have significant implications for cancer therapy, particularly in its potential role in modulating tumor progression and cellular response to treatment. Together with reported fentanyl-induced JNK activation via ROS in lung cancer models, our findings indicate that fentanyl broadly perturbs MAPK signaling networks, with divergent effects on cell fate determined by dose and redox status.

The Ramanomics evaluation of fentanyl-treated A549 cells provides critical insights into the biochemical mechanisms underlying fentanyl-induced cellular changes. An initial increase in glycogen concentration at a therapeutic dose (10 ng/mL) is likely driven by fentanyl-induced hypoxia, which enhances glycogen synthesis via glycogen branching enzyme activity. However, at overdose levels, this trend reverses due to singlet oxygen (^1^O_2_)-mediated glycogen breakdown, disrupting normal metabolic homeostasis. These distinct glycogen fluctuations offer a potential biomarker for differentiating overdose effects from therapeutic administration in lung cancer patients, particularly in the context of Raman biopsy-based diagnostics ^24^ (Fig. 7).

Lipid profiling using Ramanomics has emerged as a valuable tool for deciphering biochemical processes at the single-organelle level. By leveraging Raman spectroscopy, this approach enables high-resolution analysis of lipid composition, distribution, and metabolic dynamics within individual organelles. Such insights are crucial for understanding cellular lipid homeostasis, organelle-specific lipid metabolism, and pathological alterations linked to disease states^13,25^. Singlet oxygen (^1^O_2_)-mediated peroxidation of mitochondrial polyunsaturated fatty acids (PUFAs) leads to the formation of reactive -ene lipid products, including 4-hydroxynonenal, arachidonic acid, and hydroperoxy/hydroxyl eicosatetraenoic acid derivatives. This lipid oxidation is observed as an increase in mitochondrial unsaturation upon fentanyl overdose (Fig. 7d) ^26,27^. Upon fentanyl exposure at concentrations ranging from 100 to 1000 ng/mL, we speculate an enhanced conversion of cholesterol into cholesteryl esters. This phenomenon can be attributed to the high abundance of PUFAs in mitochondria, which serve as fatty acid precursors during esterification, catalyzed by Acyl-coenzyme A:cholesterol acyltransferase^28^. The resultant increase in mitochondrial hydrophobicity may facilitate lipid storage and transport within lipoproteins. Fentanyl-induced elevations in cholesteryl ester concentrations (Fig. 7c) observed in our study suggest a significant role for these lipid derivatives in mitochondrial lipid metabolism, potentially influencing cellular lipid homeostasis under drug-induced stress conditions. The post-peroxidation fate of mitochondrial lipids in A549 cells can be elucidated through subsequent mitochondrial lipid reactions. Oxidized polyunsaturated fatty acids (PUFAs) undergo further biochemical modifications, including enzymatic and non-enzymatic pathways that influence lipid remodeling, degradation, and metabolic adaptation within mitochondria.

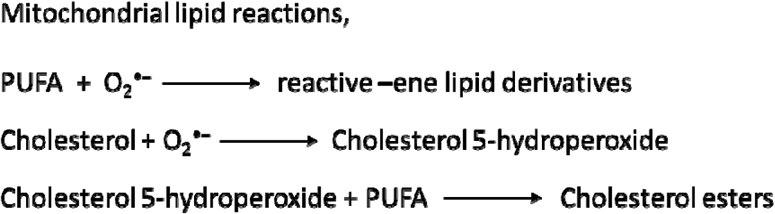

The above reactions were derived from earlier reports and our experimental observations using Ramanomics analysis^26,27^. These mitochondrial lipid processing mechanisms play a crucial role in cellular adaptation to oxidative stress and fentanyl-induced metabolic disruption, providing key insights into lipid metabolism in drug-exposed A549 cells.

Our Ramanomics data shows that fentanyl at 100-1000ng/ml induces Mitochondrial dysfunction which modulates changes at nucleic acid level as evident by the modulation in DNA, t-RNA and m-RNA (Fig 7e) at molecular levels ^29^. We also observed a Fentanyl induced elevation in accumulation of misfolded proteins generated in the mitochondrial matrix^30^ shown Fig 7f. The accumulation of misfolded proteins in the mitochondrial matrix can trigger mitophagy, a process where damaged mitochondria are selectively degraded. Fentanyl induces an elevation in misfolded protein accumulation, which suggests that mitochondrial quality control mechanisms are being overwhelmed, potentially leading to cellular stress and dysfunction and this could have implications for neurodegenerative diseases and fentanyl-related toxicity. Further, μ-opioid receptor activation can stimulate Mitogen-activated protein kinase (MAPK) activity and chronic Fentanyl exposure (100-1000 ng/mL) can trigger inhibition of MAPK pathways specifically, ERK inhibition, contributing to tolerance and dependence^31,32^. ERK activity is essential for synaptic plasticity, learning, and neuronal survival. When ERK signaling is suppressed, opioid tolerance and dependence can be exacerbated, as neurons fail to adapt effectively to fentanyl exposure, leading to compensatory changes that drive drug-seeking behavior and withdrawal symptoms. Additionally, reduction in ERK activity may impair the brain’s ability to counteract fentanyl-induced neurotoxicity, contributing to long-term cognitive and behavioral consequences. We observed an increased accumulation of phosphocholine^33^ in mitochondria (Fig 7g) and speculate that this is may be attributed to a decrease in MAPK1 which is responsible for conversion of phosphocholine to phosphotidylcholine. Additionally, Phosphocholine (precursor in phosphotidylcholine synthesis by Kennedy pathway) was also found to accumulate in vicinity of mitochondria which supports Fentanyl induced mitochondrial dysfunction ^34^. Phosphotidylcholine serves as the significant chemical backbone of both mitochondrial membrane (inner/outer) and its depletion leads to mitochondria degradation followed by cell death. Phosphocholine accumulation near mitochondria further supports our observation that fentanyl induces mitochondrial dysfunction. Since phosphocholine is a precursor for phosphatidylcholine, its buildup suggests impaired lipid metabolism, which is crucial for maintaining mitochondrial membrane integrity and function. Disruptions in phosphatidylcholine synthesis can lead to altered mitochondrial dynamics, affecting energy production and cellular homeostasis.as depicted schematically in Fig 8.

**Figure 8.**
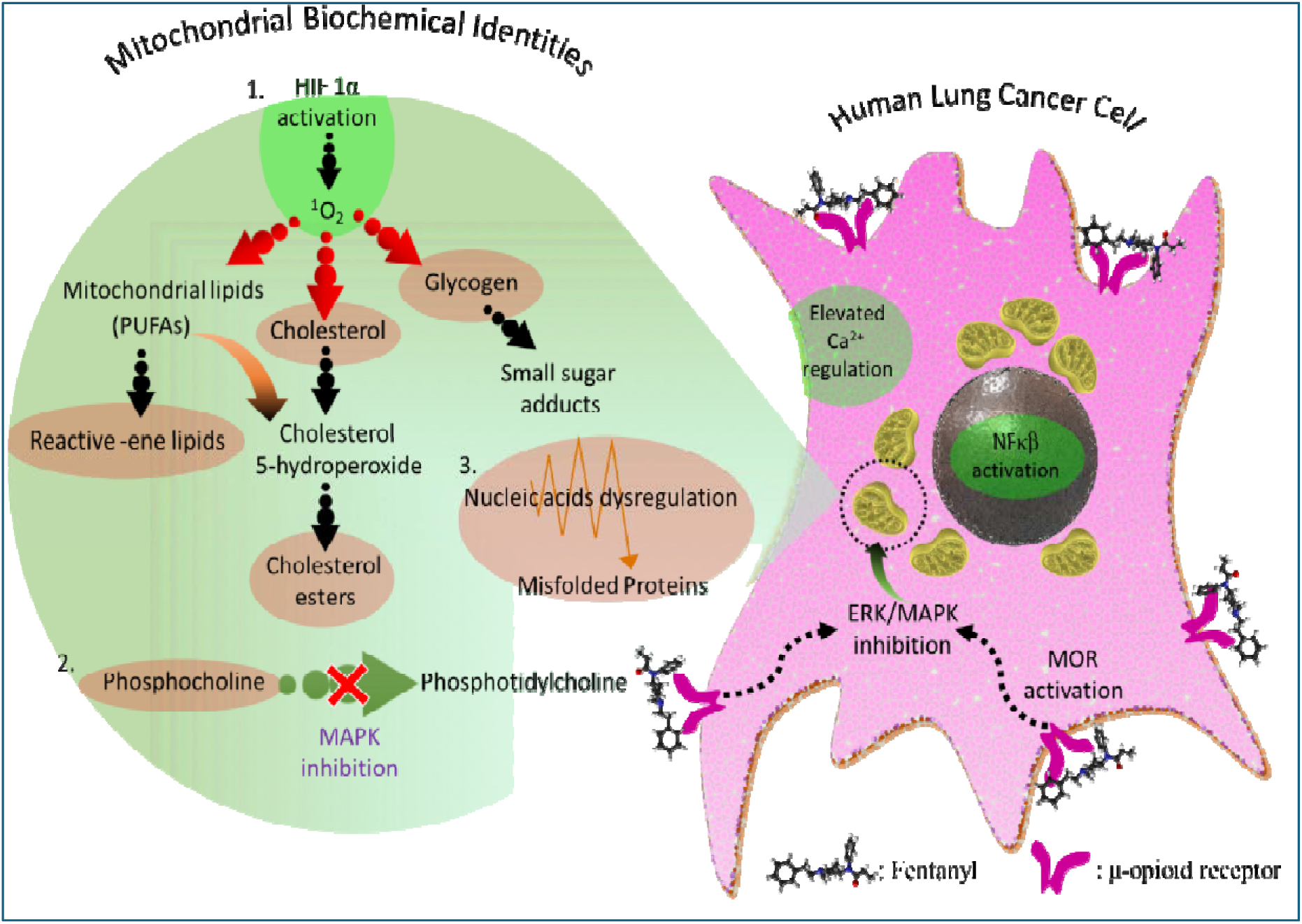
Schematic representation of cellular and mitochondrial biochemical alterations in A549 cells undergoing fentanyl-induced apoptosis. Illustration outlines stepwise biochemical events associated with chronic fentanyl exposure (100 ng/mL) in the A549 cell line, highlighting key molecular components contributing to apoptotic cell death. Cellular and mitochondrial changes are depicted with emphasis on specific pathways and molecular players. Green circular outlines denote bioassay-based observations, while orange circular outlines indicate Raman spectroscopy signatures derived from Ramanomics analysis.

## CONCLUSION

A fentanyl overdose can cause respiratory depression leading to brain hypoxia at much faster rate when compared to other opioids, therefore, understanding the biochemical effects that underlie chronic fentanyl use during lung cancer treatment is of critical importance. Our study highlights the multifaceted effects of fentanyl on A549 lung cancer cells, emphasizing its role in mitochondrial dysfunction, oxidative stress, apoptosis activation, hypoxia-inducing factor expression, and cytokine production. While fentanyl is widely used for cancer pain management, our findings suggest that it may have unintended consequences on lung cancer progression and cellular metabolism. These insights underscore the need for clinicians to exercise caution when prescribing transdermal fentanyl patches for lung cancer patients, considering its potential impact on tumor biology, due to its potential to disrupt mitochondrial function and promote oxidative stress. While some studies suggest fentanyl may suppress tumor progression, others highlight risks of immune suppression, enhanced metastasis, or mitochondrial toxicity depending on dose and context. Further research is essential to fully elucidate fentanyl’s effects and optimize its use in cancer treatment while mitigating possible risks.

## ACKNOWLEDGMENTS

The Institute for Lasers, Photonics and Biophotonics acknowledges support from the office of the Vice President for Research and Economic Development at the University at Buffalo.

